# Hebbian Wiring Plasticity Generates Efficient Network Structures for Robust Inference with Synaptic Weight Plasticity

**DOI:** 10.1101/024406

**Authors:** Naoki Hiratani, Tomoki Fukai

## Abstract

In the adult mammalian cortex, a small fraction of spines are created and eliminated every day, and the resultant synaptic connection structure is highly nonrandom, even in local circuits. However, it remains unknown whether a particular synaptic connection structure is functionally advantageous in local circuits, and why creation and elimination of synaptic connections is necessary in addition to rich synaptic weight plasticity. To answer these questions, we studied an inference task model through theoretical and numerical analyses. We demonstrate that a robustly beneficial network structure naturally emerges by combining Hebbian-type synaptic weight plasticity and wiring plasticity. Especially in a sparsely connected network, wiring plasticity achieves reliable computation by enabling efficient information transmission. Furthermore, the proposed rule reproduces experimental observed correlation between spine dynamics and task performance.

## Introduction

The amplitude of excitatory and inhibitory postsynaptic potentials (EPSPs and IPSPs), often referred to as synaptic weight, is considered a fundamental variable in neural computation(Bliss and Collingridge, 1993)(Dayan and Abbott, 2005). In the mammalian cortex, excitatory synapses often show large variations in EPSP amplitudes(Song et al., 2005)(Ikegaya et al., 2013)(Buzsáki and Mizuseki, 2014), and the amplitude of a synapse can be stable over trials(Lefort et al., 2009) and time(Yasumatsu et al., 2008), enabling rich information capacity compared with that at binary synapses(Brunel et al., 2004)(Hiratani et al., 2013). In addition, synaptic weight shows a wide variety of plasticity which depend primarily on the activity of presynaptic and postsynaptic neurons(Caporale and Dan, 2008)(Feldman, 2009). Correspondingly, previous theoretical results suggest that under appropriate synaptic plasticity, a randomly connected network is computationally sufficient for various tasks(Maass et al., 2002)(Ganguli and Sompolinsky, 2012).

On the other hand, it is also known that synaptic wiring plasticity and the resultant synaptic connection structure are crucial for computation in the brain(Chklovskii et al., 2004)(Holtmaat and Svoboda, 2009). Elimination and creation of dendritic spines are active even in the brain of adult mammalians. In rodents, the spine turnover rate is up to 15% per day in sensory cortex(Holtmaat et al., 2005) and 5% per day in motor cortex(Zuo et al., 2005). Recent studies further revealed that spine dynamics are tightly correlated with the performance of motor-related tasks(Yang et al., 2009)(Xu et al., 2009). Previous modeling studies suggest that wiring plasticity helps memory storage (Poirazi and Mel, 2001)(Stepanyants et al., 2002)(Knoblauch et al., 2010). However, in those studies, EPSP amplitude was often assumed to be a binary variable, and wiring plasticity was performed in a heuristic manner. Thus it remains unknown what should be encoded by synaptic connection structure when synaptic weights have a rich capacity for representation, and how such a connection structure can be achieved through a local spine elimination and creation mechanism, which is arguably noisy and stochastic (Kasai et al., 2010).

To answer these questions, we constructed a theoretical model of an inference task. We first studied how sparse connectivity affects the performance of the network by analytic consideration and information theoretic evaluations. Then, we investigated how synaptic weights and connectivity should be organized to perform robust inference, especially under the presence of variability in the input structure. Based on these insights, we proposed a local unsupervised rule for wiring and synaptic weight plasticity. In addition, we demonstrated that connection structure and synaptic weight learn different components under a dynamic environment, enabling robust computation. Lastly, we investigated whether the model is consistent with various experimental results on spine dynamics.

## Results

### Connection structure reduces signal variability in sparsely connected networks

What should be represented by synaptic connections and their weights, and how are those representations acquired? To explore the answers to these questions, we studied a hidden variable estimation task (**Fig. 1A**), which appears in various stages of neural information processing(Beck et al., 2008)(Lochmann and Deneve, 2011). In the task, at every time *t,* one hidden state is sampled with equal probability from *p* number of external states *s^t^* = {*0,1,…,p-1*}. Neurons in the input layer show independent stochastic responses 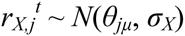 due to various noises (**Fig. 1B** middle), where *θ_jμ_* is the average firing rate of neuron *j* to the stimulus *μ*, and *σ_X_* is the constant noise amplitude. Although, we used Gaussian noise for analytical purposes, the following argument is applicable for any stochastic response that follows a general exponential family, including Poisson firing (**Supplementary Fig. 1**). Neurons in the output layer estimate the hidden variable from input neuron activity and represent the variable with population firing {*r_Y,i_*}. This task is computationally difficult because most input neurons have mixed selectivity for several hidden inputs, and the responses of the input neurons are highly stochastic (**Fig. 1C**). Let us assume that the dynamics of output neurons are written as follows:

**Figure 1:**
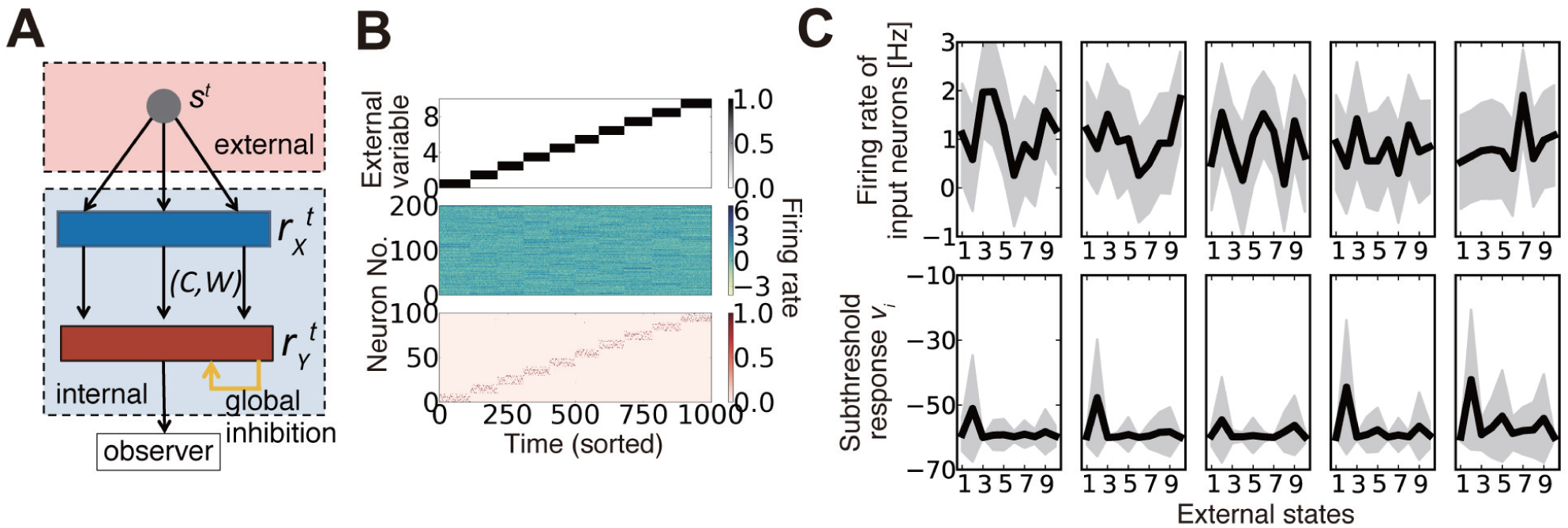
Description of the model. (**A**) Schematic diagram of the model. (**B**) An example of model behavior calculated at *ρ* = 0.16, when the synaptic connection is organized using the weight-coding scheme. The top panel represents the external variable, which takes an integer 0 to 9 in the simulation. The middle panel is the response of input neurons, and the bottom panel shows the activity of output neurons. In the simulation, each external state was randomly presented, but here the trials are sorted in ascending order. (**C**) Examples of neural activity in a simulation. Graphs on the top row represent the average firing rates of five randomly sampled input neurons for given external states (black lines) and their standard deviation (gray shadows). The bottom graphs are subthreshold responses of output neurons that represent the external state *s* = 1. Because the boundary condition for the membrane parameter 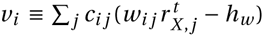 was introduced as *v_i_* > max*_i_*_′_ {*v_i_*_′_-*v_d_*}, *v_i_* is typically bounded at *–v_d_*. Note that *v_i_* is the unnormalized log-likelihood, and the units on the y-axis are arbitrary.

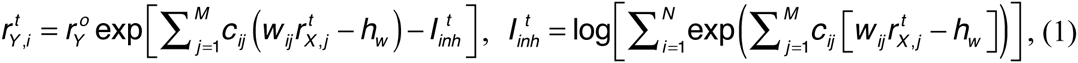

where *c_ij_* (= 0 or 1) represents connectivity from input neuron *j* to output neuron *i, w_ij_* is its synaptic weight (EPSP size), and *h_w_* is the threshold. *M* and *N* are population sizes of the input and output layers, respectively. In the model, all feedforward connections are excitatory, and the inhibitory input is provided as the global inhibition *I_inh_^t^.*

If the feedforward connection is all-to-all (i.e., *c_ij_* = 1 for all *i,j* pairs), by setting the weights as 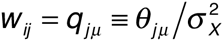 for output neuron *i* that represents external state *μ*, the network gives an optimal inference from the given firing rate vector 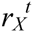, because the value *q_jμ_* represents how much evidence the firing rate of neuron *j* provides for a particular external state *μ*. (For details, see *Methods 1.1*). However, if the connectivity between the two layers is sparse, as in most regions of the brain(Potjans and Diesmann, 2014), optimal inference is generally unattainable because each output neuron can obtain a limited set of information from the input layer. How should one choose connection structure and synaptic weights in such a case? Intuitively, we could expect that if we randomly eliminate connections while keeping the synaptic weights of output neuron *i* that represents external state *μ* as *w_ij_* ∝ *q_jμ_* (below, we call it as weight coding), the network still works at a near-optimal accuracy. On the other hand, even if the synaptic weight is a constant value, if the connection probability is kept at *ρ_ij_* ∝ *q_jμ_* (i.e. connectivity coding; see *Methods 1.2* for details of coding strategies), the network is expected to achieve near-optimal performance. **Figure 2A** describes the connection matrices between input/output layers in two strategies. In the weight coding, if we sort input neurons with their preferred external states, the diagonal components of the connection matrix show high synaptic weights, whereas in the connectivity coding, the diagonal components show dense connection (**Fig. 2A**). Both of realizations asymptotically converge to optimal solution when the number of neurons in the middle layer is sufficiently large, though in a finite network, not strictly optimal under given constraints. In addition, both of them are obtainable through biologically plausible local Hebbian learning rules as we demonstrate in subsequent sections.

**Figure 2:**
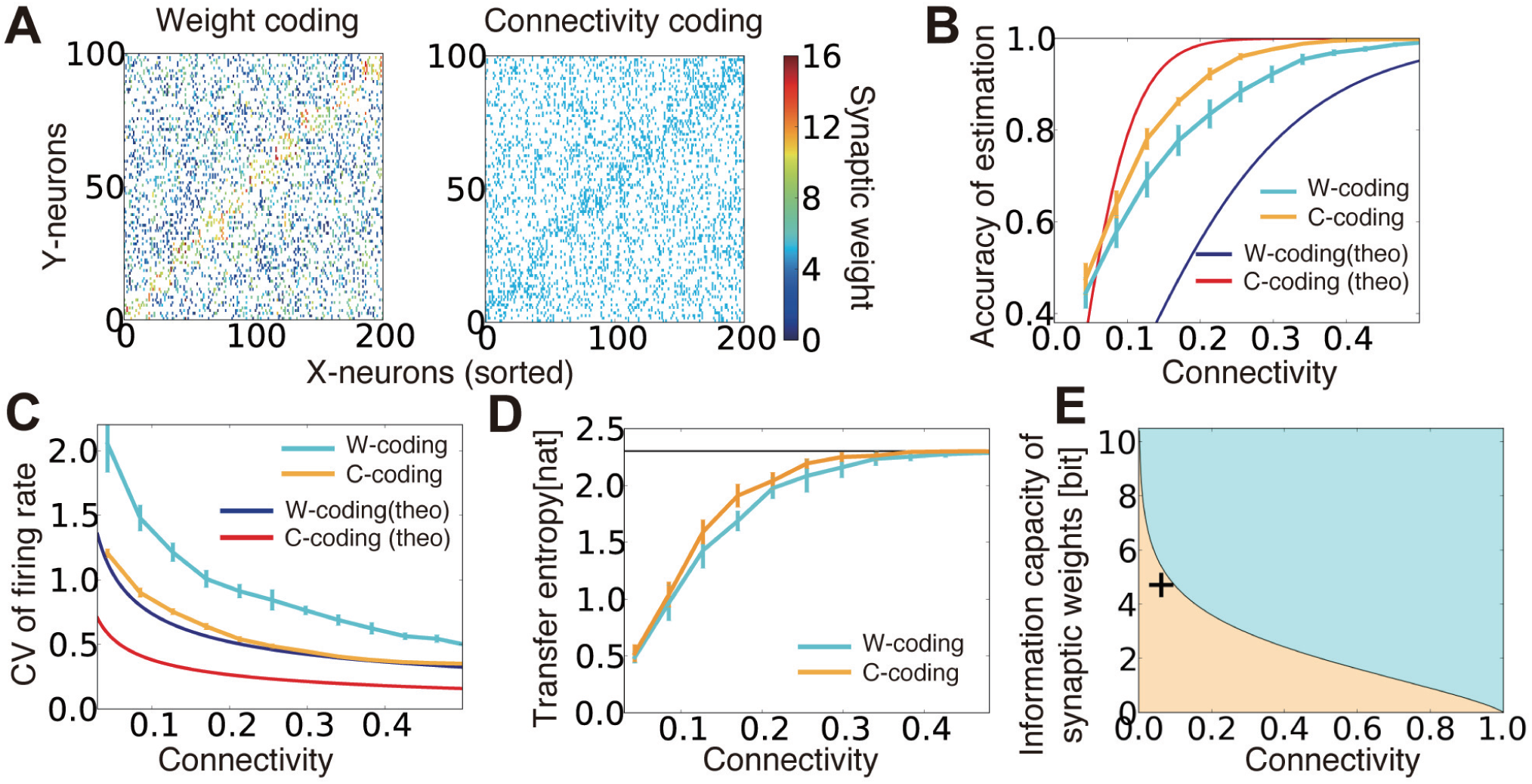
Performance comparison between connectivity coding and weight coding. (**A**) Examples of synaptic weight matrices in weight coding (W-coding) and connectivity coding (C-coding) schemes. X-neurons were sorted by their selectivity for external states. (**B**) Comparison of the performance between connectivity coding and weight coding schemes at various sparseness of connectivity. Orange and cyan lines are simulation results. The error bars represent standard deviation over 10 independent simulations. In the following panels, error bars are trial variability over 10 simulations. Red and blue lines are analytical results. (**C**) Analytically evaluated coefficient of variation (CV) of output firing rate and corresponding simulation results. For simulation results, the variance was evaluated over whole output neurons from their firing rates for their selective external states. (**D**) Estimated maximum transfer entropy for two coding strategies. Black horizontal line is the maximal information *log_e_p*. (**E**) Relative information capacity of connection structure versus synaptic weight is shown at various values of synaptic connectivity. In the orange (cyan) area, the synaptic connectivity has higher (lower) information capacity than the synaptic weights. Plus symbol represents the data point obtained from CA3-to-CA1 connections.

We evaluated the accuracy of the external state estimation using a bootstrap method (*Methods 3.2*) for both coding strategies. Under intermediate connectivity, both strategies showed reasonably good performance (as in **Fig. 1B** bottom). Intriguingly, in sparsely connected networks, the connectivity coding outperformed the weight coding, despite its binary representation (**Fig. 2B** cyan/orange lines). The analytical results confirmed this tendency (**Fig. 2B** red/blue lines; see *Methods 2.1* for the details) and indicated that the firing rates of output neurons selective for the given external state show less variability in connectivity coding than in the weight coding, enabling more reliable information transmission (**Fig. 2C**). To further understand this phenomenon, we evaluated the maximum transfer entropy of the feed forward connections: 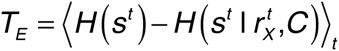. Because of limited connectivity, each output neuron obtains information only from the connected input neurons. Thus, the transfer entropy was typically lower under sparse than under dense connections in both strategies (**Fig. 2D**). However, in the connectivity coding scheme, because each output neuron can get information from relevant input neurons, the transfer entropy became relatively large compared to the weight coding (orange line in **Fig. 2D**). Therefore, analyses from both statistical and information theory-based perspectives confirm the advantage of connectivity coding over the weight coding in the sparse regions.

The result above can also be extended to arbitrary feedforward network as below. For a feedforward network of *M* times *N* neurons with connection probability *ρ*, information capacity of connections is given as *I_C_* (*ρ*) ≡ log *_MN_C_ρMN_* ≈ *MN · H*(*ρ*), where *H* represents the entropy function *H*(*ρ*) ≡ –*ρ*log*ρ*–(1–*ρ*)log(1–*ρ*). Similarly, for a given connections between two layers, information capacity of synaptic weights is written as *I_w_* (*ρ*) ≡ *ρMN*log*b*, where *b* is the number of distinctive synaptic states (Varshney et al., 2006). Therefore, when the connection probability *ρ* satisfies 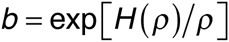, synaptic connections and weights have the same information capacities. This means that, as depicted in **Figure 2E**, in a sparsely connected network, synaptic connections tend to have larger relative information capacity, compared to a dense network with the same *b*. This result is consistent with the model above, because stochastic firing of presynaptic neuron can be translated as synaptic noise. Furthermore, in the CA3-to-CA1 connection of mice, connection probability is estimated to be around 6% (Sayer et al., 1990), and information capacity of synaptic weight is around 4.7 bits (Bartol et al., 2015), thus the connection structure should also play an active role in neural coding in the real brain (data point in **Fig. 2E**).

### Dual coding by synaptic weights and connections enables robust inference

In the section above, we demonstrated that a random connection structure highly degrades information transmission in a sparse regime to the degree that weight coding with random connection fell behind connectivity coding with a fixed weight. Therefore, in a sparse regime, it is necessary to integrate representations by synaptic weights and connections, but how should we achieve such a representation? Theoretically speaking, we should choose a connection structure that minimizes the loss of information due to sparse connectivity. This can be achieved by minimizing the KL-divergence between the distribution of the external states estimated from the all-to-all network, and the distribution estimated from a given connection structure (i.e. 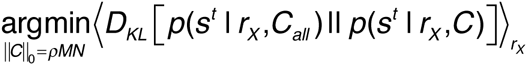, see *Methods 2.2* for details). However, this calculation requires combinatorial optimization, and local approximation is generally difficult(Donoho, 2006), thus expectedly the brain employs some heuristic alternatives. Experimental results indicate that synaptic connections and weights are often representing similar features. For example, the EPSP size of a connection in a clustered network is typically larger than the average EPSP size(Lefort et al., 2009)(Perin et al., 2011), and a similar property is suggested to hold for interlayer connections(Yoshimura et al., 2005) (Ryan et al., 2015). Therefore, we could expect that by simply combining the weight coding and connectivity coding in the previous section, low performance at the sparse regime can be avoided. On the other hand, in the previous modeling studies, synaptic rewiring and resultant connection structure were often generated by cut-off algorithm in which a synapse is eliminated if the weight is smaller than the given criteria (Chechik et al., 1998)(Navlakha et al., 2015). Thus, let us next compare the representation by combining the weight coding and connectivity coding (we call it as the dual coding below), with the cut-off coding strategy.

**Figure 3A** describes the synaptic weight distributions in the two strategies, as well as in random connection (see *Methods 1.3* for details of the implementation). When connectivity coding and weight coding are combined (i.e. in the dual coding), connection probability becomes larger in proportion to its synaptic weight (**Fig. 3A** middle), and the resultant distribution exhibits a broad distribution as observed in the experiments (Song et al., 2005)(Ikegaya et al., 2013), whereas in the cut-off strategy, the weight distribution is concentrated at a non-zero value (**Fig. 3A** right). Intuitively, the cut-off strategy seems more selective and beneficial for inference. Indeed, in the original task, the cut-off strategy enabled near-optimal performance, though the dual coding also improved the performance compared to a randomly connected network(**Fig. 3C**). However, under the presence of variability in the input layer, cut-off strategy is no longer advantageous. For instance, let us consider the case when noise amplitude *σ_X_* is not constant but pre-neuron dependent. If the firing rate variability of input neuron *j* is given by 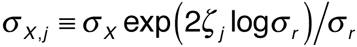, where *ζ_j_* is a random variable uniformly sampled from [0, 1), and *σ_r_* is the degree of variability, in an all-to-all network, optimal inference is still achieved by setting synaptic weights as 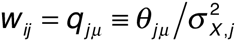. On the contrary, in the sparse region, the performance is disrupted especially in the cut-off strategy, so that the dual coding outperformed the cut-off strategy (**Fig. 3D**).

**Figure 3:**
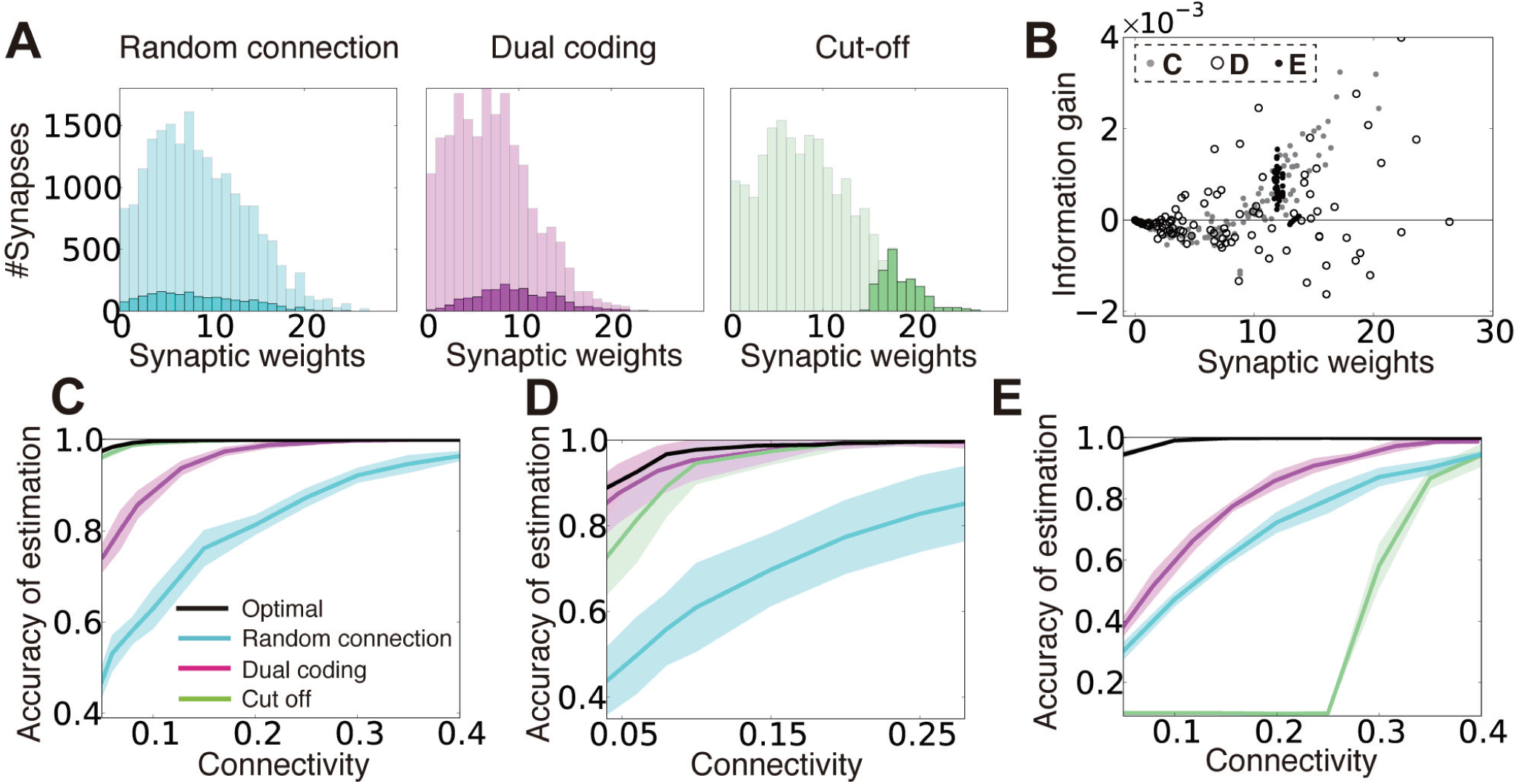
Dual coding yields robust information representation compared to fixed random connections and cut-off strategy. (**A**) Synaptic weight distributions in random connection (left), dual coding (middle), and cut-off (right) strategies. Light colors represent possible connections (i.e. distributions of synaptic weights under all-to-all connections), while dark colors show the actual connections. Connection probability was set at *ρ* = 0.1. (**B**) Relationships between the synaptic weight and the information gain per connection for three input configurations described in panels **C-E**. The open black circles were calculated with *σ_r_* = 2.0 instead of *σ_r_* = 4.0 for illustration purpose. (**C-E**) Comparisons of performance among different connection structure organizations. Note that black lines represent lower bounds for the optimal performance, but not the exact optimal solutions. In panel **D**, the means and standard deviations were calculated over 100 simulation trials instead of 10 due to intrinsic variability.

To further illustrate this phenomenon, let us next consider a case when a quarter of input neurons show a constant high response for all of the external states as 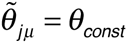, and the rest of input neurons show high response for randomly selected half of external states (i.e. 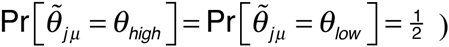), where *θ_low_ < θ_high_ < θ_const_*, and 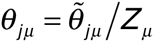 with the normalization factor 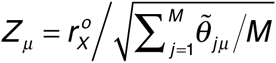. Even in this case, 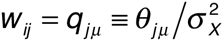 is the optimal synaptic weights configuration in the all-to-all network, but if we create a sparse network with cut-off algorithm, the performance drops dramatically at certain connectivity, whereas in the dual coding, the accuracy is kept at some high levels even in the sparse connectivity (**Fig. 3E**).

To get insights on why the dual coding is more robust against variability in the input layer, for three input configurations described above, we calculated the relationship between synaptic weight *w_ij_* and the information gained by a single synaptic connection *ΔI_ij_.* Here, we defined the information gain *ΔI_ij_* by the mean reduction in the KL divergence 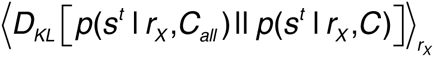, achieved by adding one synaptic connection *c_ij_* to a randomly connected network *C* (see *Method 2.2* for details). In the original model, *ΔI_ij_* has nearly a linear relationship with the synaptic weight *w_ij_* (gray points in **Fig. 3B**), thus by simply removing the connections with small synaptic weights, a near-optimal connection structure was acquired (**Fig. 3C**). On the other hand, when the input layer is not homogeneous, large synapses tend to have negative (black circles in **Fig. 3B**) or zero (black points in **Fig. 3B**) gains, as a result, the linear relationship between the weight and the information gain was lost. Thus, in these cases, the dual coding is less likely to be disrupted by non-beneficial connections.

Although our consideration here is limited to a specific realization of synaptic weights, in general, it is difficult to represent the information gain by locally acquired synaptic weight, so we could expect that the cut-off strategy is not the optimal connectivity organization in many cases.

### Local Hebbian learning of the dual coding

The argument in the previous section suggest that, by combining the weight coding and connectivity coding, the network can robustly perform inference especially in sparsely connected regions. However, in the previous sections, a specific connection and weight structure were given a priori, although structures in local neural circuits are expected to be obtained with local weight plasticity and wiring plasticity. Thus, we next investigate whether dual coding can be achieved through a local unsupervised synaptic plasticity rule.

Let us first consider learning of synaptic weights. In order to achieve the weight coding, synaptic weight *w_ij_* should converge to 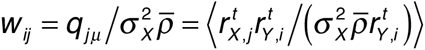 when output neuron *i* represents external state *μ*, and 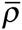 represents the mean connectivity of the network. Thus, synaptic weight change 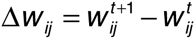 is given as:

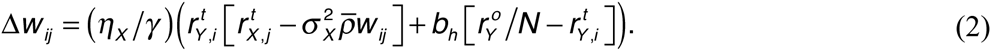

The second term is the homeostatic term heuristically added to constrain the average firing rates of output neurons(Turrigiano and Nelson, 2004). Note that the first term corresponds to stochastic gradient descending on 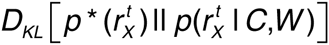, because the weight coding approximates the optimal representation by synaptic weights (Nessler et al., 2013)(see *Methods 1.4* for details). We performed this unsupervised synaptic weight learning on a randomly connected network. When the connectivity is sufficiently dense, the network successfully acquired a suitable representation (**Fig. 4A**). Especially under a sufficient level of homeostatic plasticity (**Fig. 4B**), the average firing rate showed a narrow unimodal distribution (**Fig. 4C** top), and most of the output neurons acquired selectivity for one of external states (**Fig. 4C** bottom).

**Figure 4:**
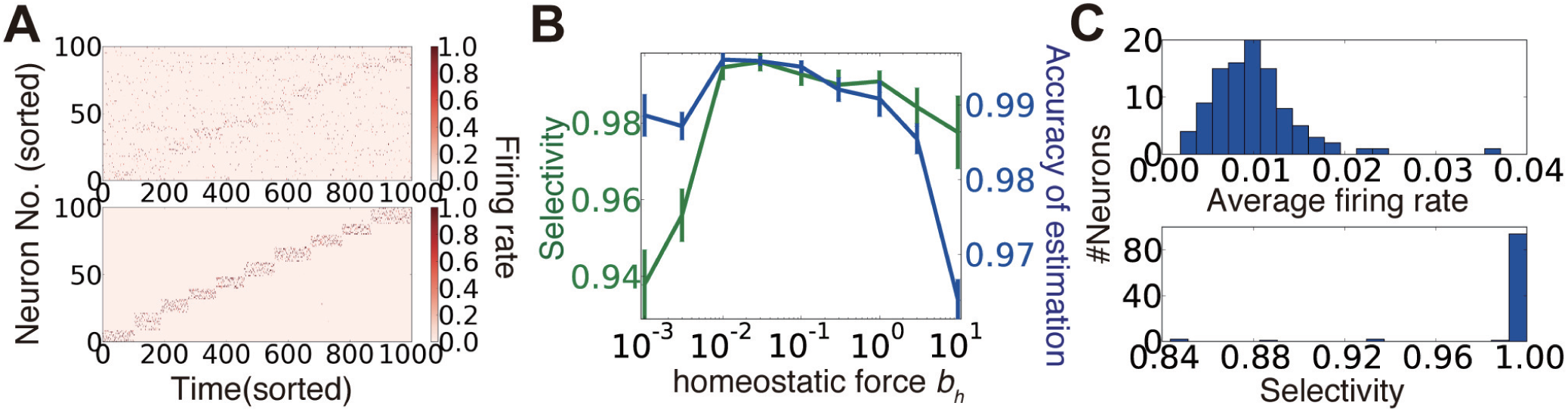
Synaptic weight learning on random connection structures. (**A**) An example of output neuron activity before (top) and after (bottom) synaptic weight learning calculated at connectivity *ρ* = 0.4. (**B**) Selectivity of output neurons and accuracy of estimation at various strengths of homeostatic plasticity at *ρ* = 0.4. Selectivity was defined as 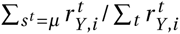 for *i* ∈ Ω*_μ_.* (**C**) Histogram of average firing rates of output neurons (top), and selectivity of each neuron calculated for the simulation depicted in panel **A**.

We next investigated the learning of connection structures by wiring plasticity. Unlike synaptic weight plasticity, it is not yet well understood how we can achieve functional connection structure with local wiring plasticity. In particular, rapid rewiring may disrupt the network structure, and possibly worsen the performance (Chechik et al., 1998). Thus, let us first consider a simple rewiring rule, and discuss the biological correspondence later. Here, we introduced a variable *ρ_ij_*, for each combination (i,j) of presynaptic neuron *j* and postsynaptic neuron *i*, which represents the connection probability. If we randomly create a synaptic connection between neuron (i,j) with probability *ρ_ij_/τ_c_* and eliminate it with probability (1- *ρ_ij_*)/*τ_c_*, on average there is a connection between neuron (i,j) with probability *ρ_ij_,* when the maximum number of synaptic connections is bounded by 1. In this way, the total number of synaptic connections is kept constant on average, without any global regulation mechanism.

From a similar argument done for synaptic weights, the learning rule for connection probability *ρ_ij_* is derived as:

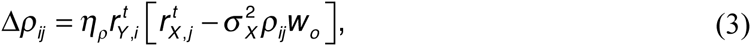

where *w_o_* is the expected mean synaptic weight (*Methods 1.5*). Under this rule, the connection probabilities converge to the connectivity coding. Moreover, although this rule does not maximize the transfer entropy of the connections, direction of learning is on average close to the direction of the stochastic gradient on transfer entropy. Therefore, the above rule does not reduce the transfer entropy of the connection on average (see *Methods 1.6).*

**Figure 5A** shows the typical behavior of *ρ_ij_* and *w_ij_* under combination of this wiring rule (equation (3)) and the weight plasticity rule described in equation (2) (we call this combination as the dual Hebbian rule because both equations (2) and (3) have Hebbian forms). When the connection probability is low, connections between two neurons are rare, and, even when a spine is created due to probabilistic creation, the spine is rapidly eliminated (**Fig. 5A** top). In the moderate connection probability, spine creation is more frequent, and the created spine survives longer (**Fig. 5A** middle). When the connection probability is high enough, there is almost always a connection between two neurons, and the synaptic weight of the connection is large because synaptic weight dynamics also follow a similar Hebbian rule (**Fig. 5A** bottom).

**Figure 5:**
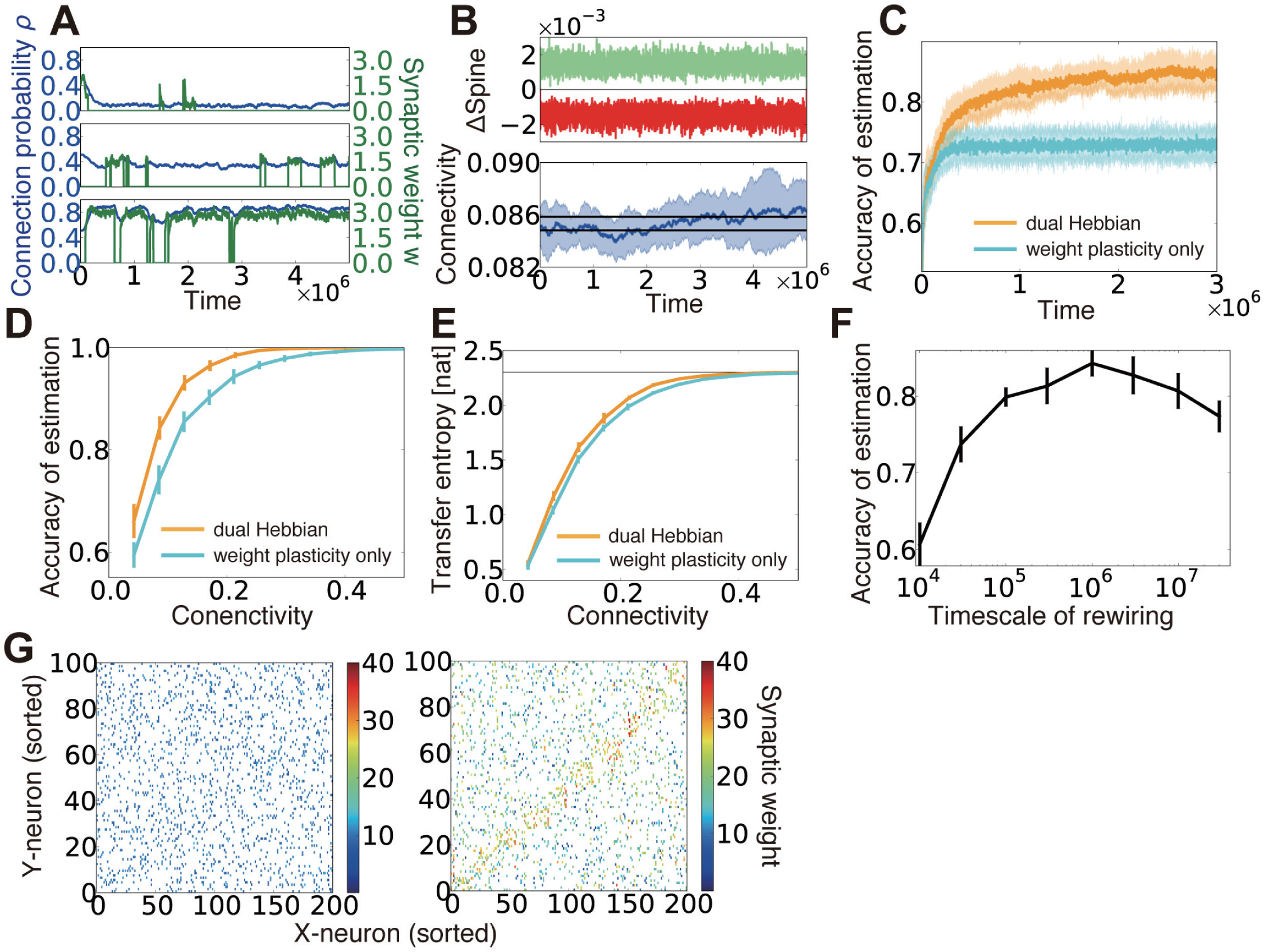
Dual Hebbian learning for synaptic weights and connections. (**A**) Examples of spine creation and elimination. In all three panels, green lines show synaptic weights, and blue lines are connection probability. When there is not a synaptic connection between two neurons, the synaptic weight becomes zero, but the connection probability can take a non-zero value. Simulation was calculated at *ρ* = 0.48, *η_ρ_* = 0.001, and *τ_c_* = 10^5^. (**B**) Change in connectivity due to synaptic elimination and creation. Number of spines eliminated (red) and created (green) per unit time was balanced (top). As a result, connectivity did not appreciably change due to rewiring (bottom). Black lines in the bottom graph are the mean connectivity at *γ* = 0.1 and *γ* = 0.101 in the model without rewiring. (**C**) Accuracy of estimation for the model with/without wiring plasticity. For the dual Hebbian model, the sparseness parameter was set as *γ* = 0.1, whereas *γ* = 0.101 was used for the weight plasticity model to perform comparisons at the same connectivity (see panel **B**). (**D, E**) Comparison of the performance (**D**) and the maximum estimated transfer entropy (**E**) after learning between the dual Hebbian model and the model implemented with synaptic plasticity only at various degrees of connectivity. Horizontal line in panel **E** represents the total information *log_e_p*. (**F**) Accuracy of estimation with various timescales for rewiring *τ_c_*. Note that the simulation was performed only for 5 × 10^6^ time steps, and the performance did not converge for the model with a longer timescale. (**G**) Synaptic weight matrices before (left) and after (right) learning. Both X-neurons (input neuron) and Y-neurons (output neurons) were sorted based on their preferred external states.

We implemented the dual Hebbian rule in our model and compared the performance of the model with that of synaptic weight plasticity on a fixed random synaptic connection structure. Because spine creation and elimination are naturally balanced in the proposed rule (**Fig. 5B** top), the total number of synaptic connections was nearly unchanged throughout the learning process (**Fig. 5B** bottom). As expected, the dual Hebbian rule yielded better performance (**Fig. 5C,D**) and higher estimated transfer entropy than the corresponding weight plasticity only model (**Fig. 5E**). This improvement was particularly significant when the frequency of rewiring was in an intermediate range (**Fig. 5F**). When rewiring was too slow, the model showed essentially the same behavior as that in the weight plasticity only model, whereas excessively frequent probabilistic rewiring disturbed the connection structure. Although a direct comparison with experimental results is difficult, the optimal rewiring timescale occurred within hours to days, under the assumption that firing rate dynamics (equation (1)) are updated every 10 to 100 ms. Initially, both connectivity and weights were random (**Fig. 5G** left), but after the learning process, the diagonal components of the weight matrix developed relatively larger synaptic weights, and, at the same time, denser connectivity than the off-diagonal components (**Fig. 5G** right). Thus, through dual Hebbian learning, the network can indeed acquire a connection structure that enables efficient information transmission between two layers; as a result, the performance improves when the connectivity is moderately sparse (**Fig. 5D, E**). Although the performance was slightly worse than that of a fully-connected network, synaptic transmission consumes a large amount of energy(Sengupta et al., 2013), and synaptic connection is a major source of noise(Faisal et al., 2008). Therefore, it is beneficial for the brain to achieve a similar level of performance using a network with fewer connections.

### Connection structure can acquire constant components of stimuli and enable rapid learning

We have shown that the dual coding by synaptic weights and connections robustly helps computation in a sparsely connected network, and the desirable weight and connectivity structures are naturally acquired through the dual Hebbian rule. Although we were primary focused on sparse regions, the rule potentially provides some beneficial effects even in densely connected networks. To consider this issue, we extended the previous static external model to a dynamic one, in which at every interval *T_2_,* response probabilities of input neurons partly change. If we define the constant component as *θ_const_* and the variable component as *θ_var_,* then the total model becomes 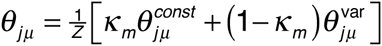, where the normalization term is given as 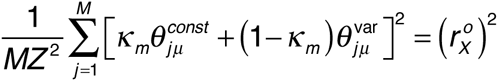 (**Fig. 6A**). In this case, when the learning was performed only with synaptic weights based on fixed random connections, although the performance rapidly improved, every time a part of the model changed, the performance dropped dramatically and only gradually returned to a higher level (cyan line in **Fig. 6B**). By contrast, under the dual Hebbian learning rule, the performance immediately after the model shift (i.e., the performance at the trough of the oscillation) gradually increased, and convergence became faster (**Fig. 6B,C**), although the total connectivity stayed nearly the same (**Fig. 6D**). After learning, the synaptic connection structure showed a higher correlation with the constant component than with the variable component (**Fig. 6E**; see *Methods 3.3*). By contrast, at every session, synaptic weight structure learned the variable component better than it learned the constant component (**Fig. 6F**). The timescale for synaptic rewiring needed to be long enough to be comparable with the timescale of the external variability *T2* to capture the constant component. Otherwise, connectivity was also strongly modulated by the variable component of the external model (**Fig. 6G**). After sufficient learning, the synaptic weight *w* and the corresponding connection probability *ρ* roughly followed a linear relationship (**Fig. 6H**). Remarkably, some synapses developed connection probability *ρ* = 1, meaning that these synapses were almost permanently stable because the elimination probability (1-*ρ*)/*τ_c_* became nearly zero.

**Figure 6:**
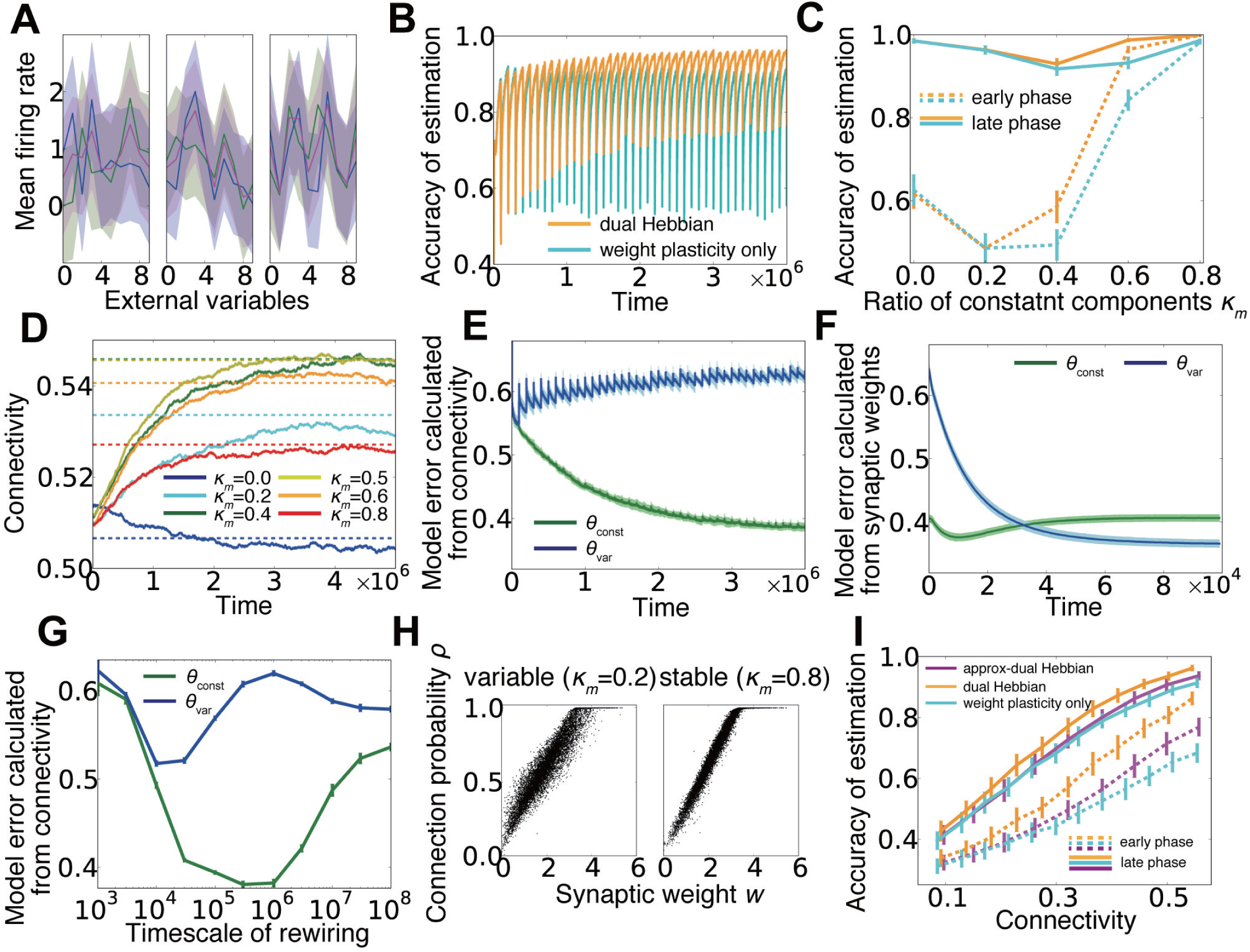
Dual learning under a dynamic environment. (**A**) Examples of input neuron responses. Blue lines represent the constant components *θ_const_*, green lines show the variable components *θ_var_*, and magenta lines are the total external models *θ* calculated from the normalized sum. (**B**) Learning curves for the model with or without wiring plasticity, when the variable components change every 10^5^ time steps. (**C**) Accuracy of estimation for various ratios of constant components. Early phase performance was calculated from the activity within 10,000 steps after the variable component shift, and the late phase performance was calculated from the activity within 10,000 steps before the shift. As in panel **B**, orange lines represent the dual Hebbian model, and cyan lines are for the model with weight plasticity only. (**D**) Trajectories of connectivity change. Connectivity tends to increase slightly during learning. Dotted lines are mean connectivity at *(κ_m_*,*γ*) = (0.0,0.595), (0.2,0.625), (0.4,0.64), (0.5,0.64), (0.6,0.635), and (0.8,0.620). In **C**, these parameters were used for the synaptic plasticity only model, whereas *γ* was fixed at *γ* = 0.6 for the dual Hebbian model. (**E,F**) Model error calculated from connectivity (**E**) and synaptic weights (**F**). Note that the timescale of **F** is the duration in which the variable component is constant, not the entire simulation (i.e. the scale of x-axis is 10^4^ not 10^6^). (**G**) Model error calculated from connectivity for various rewiring timescales *τ_c_*. For a large *τ_c_*, the learning process does not converge during the simulation. (**H**) Relationship between synaptic weight w and connection probability *ρ* at the end of learning. When the external model is stable, *w* and *ρ* have a more linear relationship than that for the variable case. (**I**) Comparison of performances among the model without wiring plasticity (cyan), the dual Hebbian model (orange), the approximated model (magenta).

### Approximated dual Hebbian learning rule reconciles with experimentally observed spine dynamics

Our results up to this point have revealed functional advantages of dual Hebbian learning. In this last section, we investigated the correspondence between the experimentally observed spine dynamics and the proposed rule. To this end, we first studied whether a realistic spine dynamics rule approximates the proposed rule, and then examined if the rule explains the experimentally known relationship between synaptic rewiring and motor learning (Yang et al., 2009)(Xu et al., 2009).

Previous experimental results suggest that a small spine is more likely to be eliminated(Yasumatsu et al., 2008)(Kasai et al., 2010), and spine size often increases or decreases in response to LTP or LTD respectively, with a certain delay (Matsuzaki et al., 2004)(Wiegert and Oertner, 2013). In addition, though spine creation is to some extent influenced by postsynaptic activity (Knott et al., 2006)(Yang et al., 2014), the creation is expected to be more or less a random process (Holtmaat and Svoboda, 2009). Thus, changes in the connection probability can be described as

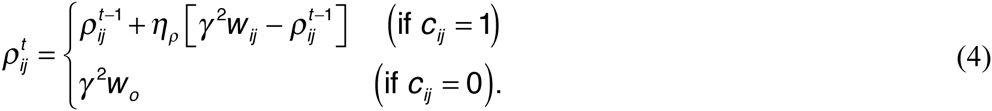

By combining this rule and the Hebbian weight plasticity described in equation (2), the dynamics of connection probability well replicated the experimentally observed spine dynamics (Yasumatsu et al., 2008)(Kasai et al., 2010) (**Fig. 7A-C**). Moreover, the rule outperformed the synaptic weight only model in the inference task, although the rule performed poorly compared to the dual Hebbian rule due to the lack of activity dependence in spine creation (magenta line in **Fig. 6I**). This result suggests that plasticity rule by equations (2) and (4) well approximates the dual Hebbian rule (equations (2)+(3)). This is because, even if the changes in the connection probability are given as a function of synaptic weight as in equation (4), as long as the weight plasticity rule follows equation (2), wiring plasticity indirectly shows a Hebbian dependency for pre- and postsynaptic activities as in the original dual Hebbian rule (equation (3)). As a result, the approximated rule gives a good approximation of the original dual Hebbian rule.

**Figure 7:**
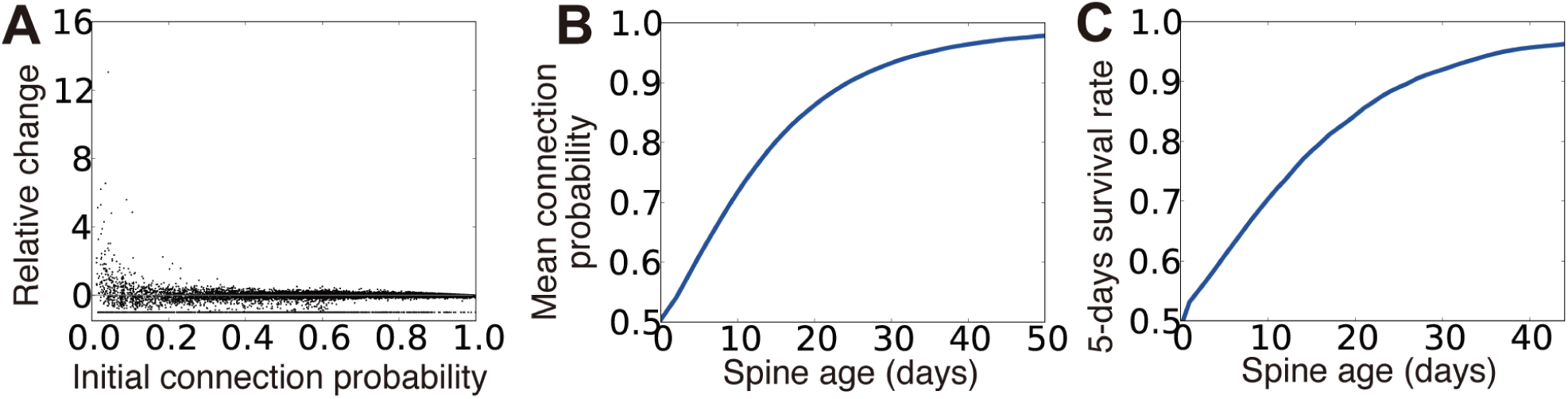
Spine dynamics of the approximated dual Hebbian model. (**A**) Relative change of connection probability in 10^5^ time steps. If the initial connection probability is low, the relative change after 10^5^ time steps has a tendency to be positive, whereas spines with a high connection probability are more likely to show negative changes. The line at the bottom represents eliminated spines (i.e., relative change = -1). (**B,C**) Relationships between spine age and the mean connection probability (**B**) and the 5-days survival rate (**C**). Consistent with the experimental results, survival rate is positively correlated with spine age. 5-days survival rate was calculated by regarding 10^5^ time steps as one day.

We next applied this approximated learning rule to motor learning tasks. The primary motor cortex has to adequately read-out motor commands based on inputs from pre-motor regions(Salinas and Romo, 1998)(Sul et al., 2011). In addition, the connection from layer 2/3 to layer 5 is considered to be a major pathway in motor learning(Masamizu et al., 2014). Thus we hypothesized that the input and output layers of our model can represent layers 2/3 and 5 in the motor cortex. We first studied the influence of training on spine survival(Xu et al., 2009) (**Fig. 8A**). To compare with experimental results, below we regarded 10^5^ time steps as one day, and described the training and control phases as two independent external models *θ_ctrl_* and *θ_train_.* In both training and control cases, newly created spines were less stable than pre-existing spines (solid lines vs. dotted lines in **Fig. 8B**), because older spines tended to have a larger connection probability (**Fig. 7B**). In addition, continuous training turned pre-existed spines less stable and new spines more stable than their respective counterparts in the control case (red lines vs. lime lines in **Fig. 8B**). The 5-day survival rate of a spine was higher for spines created within a couple of days from the beginning of training compared with spines in the control case, whereas the survival rate converged to the control level after several days of training (**Fig. 8C**). We next considered the relationship between spine dynamics and task performance(Yang et al., 2009). For this purpose, we compared task performance at the beginning of the test period among simulations with various training lengths (**Fig. 8D**). Here, we assumed that spine elimination was enhanced during continuous training, as is observed in experiments(Yang et al., 2009)(Xu et al., 2009). The performance was positively correlated with both the survival rate at day 7 of new spines formed during the first 2 days, and the elimination rate of existing spines (left and right panels of **Fig. 8E**). By contrast, the performance was independent from the total ratio of newly formed spines from day 0 to 6 (middle panel of **Fig. 8E**). These results demonstrate that complex spine dynamics are well described by the approximated dual Hebbian rule, suggesting that the brain uses a dual learning mechanism.

**Figure 8:**
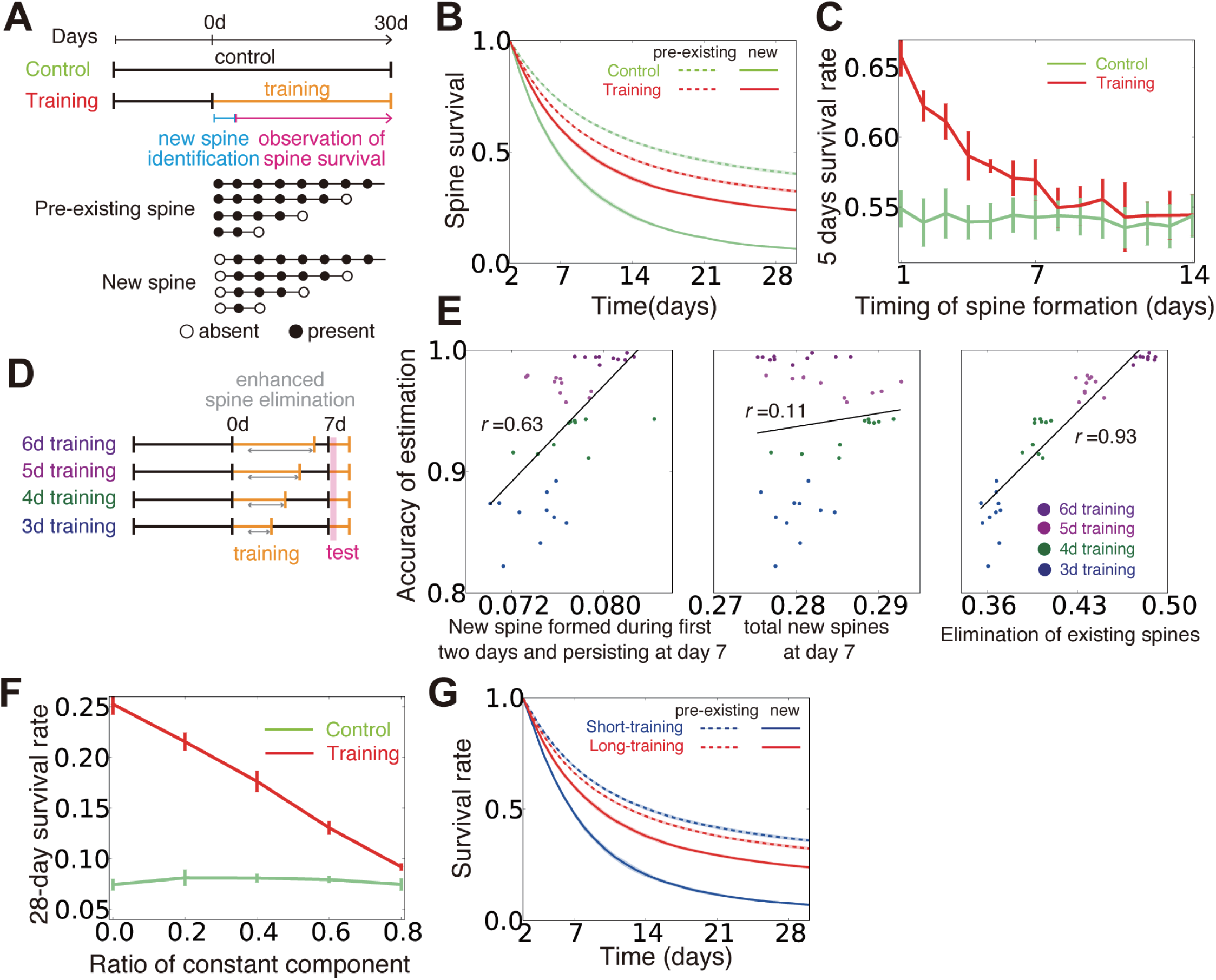
Influence of training on spine dynamics. (**A**) Schematic diagrams of the simulation protocols for panels **B,C**, and **F,G**, and examples of spine dynamics for pre-existing spines and new spines. (**B**) Spine survival rates for control and training simulations. Dotted lines represent survival rates of pre-existing spines (spines created before day 0 and existing on day 2), and solid lines are new spines created between day 0 and day 2. (**C**) The 5-day survival rate of spines created at different stages of learning. (**D,E**) Relationships between creation and elimination of spines and task performance. Performance was calculated from the activity within 2,000-7,000 time steps after the beginning of the test phase. In the simulation, the synaptic elimination was increased fivefold from day 1 to the end of training. (**F**) Effect of similarity between the control condition and training on the new spine survival rate. The value of *κ_m_* was changed as in **C** to alter the similarity between the two conditions. Note that *κ_m_* = 0 in panels **A-E**, and **G**. (**G**) Spine survival rates for short-training (2 d) and long-training (30 d) simulations. Pre-existing and new spines were defined as in panels **A,B**.

## Discussion

In this study, we first analyzed how random connection structures impair performance in sparsely connected networks by analyzing the change in signal variability and the transfer entropy in the weight coding and the connectivity coding strategies (**Fig. 2**). Subsequently, we showed that connection structures created by the cut-off strategy are not beneficial under the presence of input variability, due to lack of positive correlation between the information gain and weight of synaptic connections (**Fig. 3**). Based on these insights, we proposed that the dual coding by weight and connectivity structures as a robust representation strategy, then demonstrated that the dual coding is naturally achieved through dual Hebbian learning by synaptic weight plasticity and wiring plasticity (**Fig. 4, 5**). We also revealed that, even in a densely connected network in which synaptic weight plasticity is sufficient in terms of performance, by encoding the time-invariant components with synaptic connection structure, the network can achieve rapid learning and robust performance (**Fig. 6**). Even if spine creation is random, the proposed framework still works effectively, and the approximated model with random spine creation is indeed sufficient to reproduce various experimental results (**Fig. 7, 8**).

### Model evaluation

Spine dynamics depend on the age of the animal(Holtmaat et al., 2005), the brain region(Zuo et al., 2005), and many molecules play crucial roles(Kasai et al., 2010)(Caroni et al., 2012), making it difficult for any theoretical models to fully capture the complexity. Nevertheless, our simple mathematical model replicated many key features(Yasumatsu et al., 2008)(Yang et al., 2009)’(Xu et al., 2009)(Kasai et al., 2010). For instance, small spines often show enlargement, while large spines are more likely to show shrinkage (**Fig. 7A**). Older spines tend to have a large connection probability, which is proportional to spine size (**Fig. 7B**), and they are more stable (**Fig. 7C**). In addition, training enhances the stability of newly created spines, whereas it degrades the stability of older spines (**Fig. 8B**).

### Experimental prediction

In the developmental stage, both axon guidance(Munz et al., 2014) and dendritic extension(Matsui et al., 2013) show Hebbian-type activity dependence, but in the adult cortex, both axons and dendrites seldom change their structures(Holtmaat and Svoboda, 2009). Thus, although recent experimental results suggest some activity dependence for spine creation(Knott et al., 2006)(Yang et al., 2014), it is still unclear to what extent spine creation depends on the activity of presynaptic and postsynaptic neurons. Our model indicates that in terms of performance, spine creation should fully depend on both presynaptic and postsynaptic activity (**Fig. 6I**). However, we also showed that it is possible to replicate a wide range of experimental results on spine dynamics without activity-dependent spine creation (**Fig. 8**).

Furthermore, whether or not spine survival rate increases through training is controversial(Yang et al., 2009)(Xu et al., 2009). Our model predicts that the stability of new spines highly depends on the similarity between the new task and control behavior (**Fig. 8F**). When the similarity is low, new spines created in the new task are expected to be more stable than those created in the control case, because the synaptic connection structure would need to be reorganized. By contrast, when the similarity is high, the stability of the new spines would be comparable to that of the control. In addition, our model replicates the effect of varying training duration on spine stability(Yang et al., 2009). When training was rapidly terminated, newly formed spines became less stable than those undergoing continuous training (**Fig. 8G**).

### Related studies

Previous theoretical studies revealed candidate rules for spine creation and elimination(Deger et al., 2012)(Zheng et al., 2013)(Fauth et al., 2015), yet their functional benefits were not fully clarified in those studies. Some modeling studies considered the functional implications of synaptic rewiring (Poirazi and Mel, 2001) or optimality in regard to benefit and wiring cost (Chen et al., 2006), but the functional significance of synaptic plasticity and the variability of EPSP size were not considered in those models. In comparison, our study revealed functional roles of wiring plasticity that cooperates with synaptic weight plasticity and obeys local unsupervised rewiring rules. In addition, we extended the previous results on single-spine information storage and synaptic noise (Varshney et al., 2006) into a network, and provided a comparison with experimental results (**Fig. 2E**).

Previous studies on associative memory models found the cut-off coding as the optimal strategy for maximizing the information capacity per synapse (Chechik et al., 1998)(Knoblauch et al., 2010). Our results suggest that the above result is the outcome of the tight positive correlation between the information gain and synaptic weight in associative memory systems, and not generally applicable to other paradigms (**Fig. 3BC**). In addition, although cut-off strategy did not yield biologically plausible synaptic weight distributions in our task setting (**Fig. 3A** right), in perceptron-based models, this unrealistic situation can be avoided by tuning the threshold of neural dynamics (Brunel et al., 2004)(Sacramento et al., 2015). Especially, cut-off strategy may provide a good approximation for developmental wiring plasticity(Ko et al., 2013), though the algorithm is not fully consistent with wiring plasticity in the adult animals.

Finally, our model provides a biologically plausible interpretation for multi-timescale learning processes. It was previously shown that learning with two synaptic variables on different timescales is beneficial under a dynamically changing environment(Fusi et al., 2007). In our model, both fast and slow variables played important roles, whereas in previous studies, only one variable was usually more effective than others, depending on the task context.

## Methods

### 1. Model

#### 1.1 Model dynamics

We first define the model and the learning rule for general exponential family, and derive equations for two examples (Gaussian and Poisson). In the task, at every time *t*, one hidden state *s^t^* is sampled from prior distribution *p*(*s*). Neurons in the input layer show stochastic response 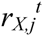 that follows probabilistic distribution *f*(*r_X,j_* | *s^t^*):

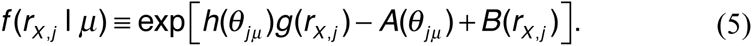

From these input neuron activities, neurons in output layer estimate the hidden variables. Here we assume maximum likelihood estimation for decision making unit, as the external state is a discrete variable. In this framework, in order to detect the hidden signal, firing rate of neuron *i* should be proportional to posterior

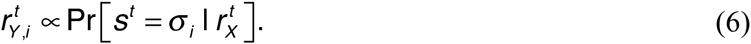

where *σ_i_* represents the index of the hidden variable preferred by output neuron *i* (Beck et al., 2008)(Lochmann and Deneve, 2011). Note that {*r_X,j_*} represent firing rates of input neurons, whereas {*r_Y,i_*} represent the rates of output neurons. Due to Bayes rule, estimation of *s^t^* is given by,

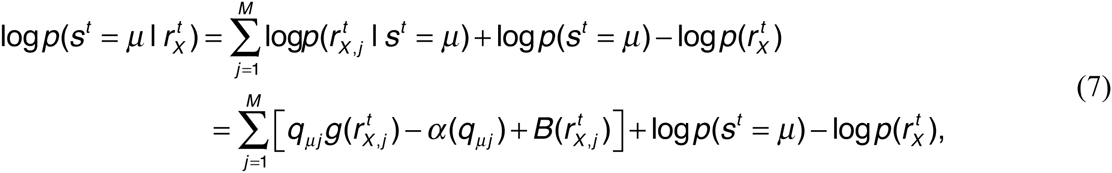

where *q_jμ_* ≡ *h*(*θ_jμ_*), 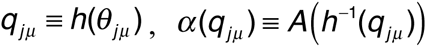. If we assume the uniformity of hidden states as log *p*(*s^t^ = μ*): *const*, and 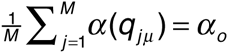, the equation above becomes

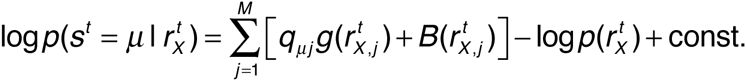

To achieve neural implementation of this inference problem, let us consider a neural dynamics in which the firing rates of output neurons follow,

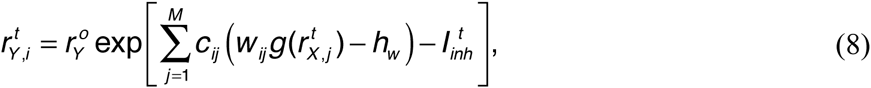

where,

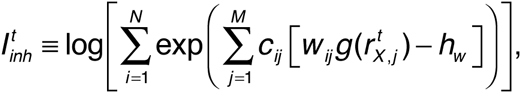

and *h_w_* is the threshold. If connection is all-to-all, *w_ij_* = *q_jμ_* gives optimal inference, because

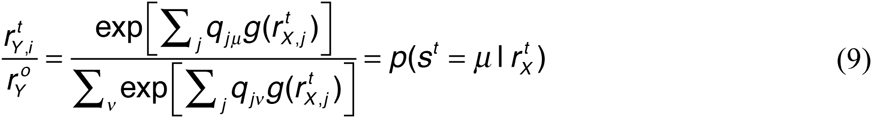

Note that *h_w_* is not necessary to achieve optimal inference, however, under a sparse connection, *h_w_* is important for reducing the effect of connection variability. In this formalization, even in non-all-to-all network, if the sparseness of connectivity stays in reasonable range, near-optimal inference can be performed for arbitrary feedforward connectivity by adjusting synaptic weight to 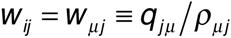 where 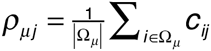.

#### 1.2. Weight coding and connectivity coding

Let us first consider the case when the connection probability is constant (i.e. *ρ_ij_*=*ρ*). By substituting *ρ_ij_*=*ρ* into the above equations, *c* and *w* are given with 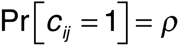 and 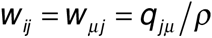, where the mean connectivity is given as 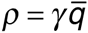, and 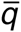 is the average of the normalized mean response *q_jμ_* (i.e., 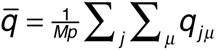). Parameter *γ* is introduced to control the sparseness of connections, and here we assumed that neuron *i* represents the external state 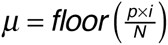 (i.e., if 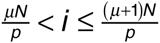, output neuron *i* represents the state *μ*). Under this configuration, the representation is solely achieved by the synaptic weights, thus we call this coding strategy as the weight coding.

On the other hand, if the synaptic weight is kept at a constant value, the representation is realized by synaptic connection structure (i.e. connectivity coding). In this case, the model is given by 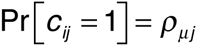 and 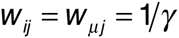, where 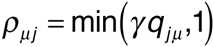.

#### 1.3 Dual coding and cut-off coding

By combining the weight coding and connectivity coding described above, the dual coding is given as *w_ij_ = w_μj_* = *q_iμ_*/*ρ*, 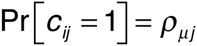, 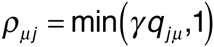, where *ρ* was defined by 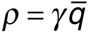, 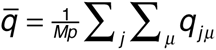, as in the weight coding. For the cut-off coding strategy, the synaptic weight was chosen as *w_ij_ = w_μj_* = *q_iμ_*/*ρ_o_* where *ρ_o_* is the mean connection probability. Based on these synaptic weights, for each output neuron, we selected *Mρ_o_* largest synaptic connections, and eliminated all other connections. Thus, connection matrix *C* was given as 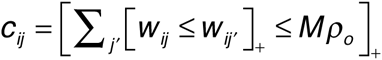, where [true]_+_ =1, [false]_+_=0. When multiple connections have the same weight, we randomly selected the connections so that the total number of inbound connections becomes *Mρ_o_*. Finally, in the random connection strategy, synaptic weights and connections were determined as *w_ij_ = w_μj_* = *q_iμ_*/*ρ_o_*, 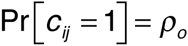.

#### 1.4 Synaptic weight learning

To perform maximum likelihood estimation from output neuron activity, synaptic weight matrix between input neurons and output neurons should provide a reverse model of input neuron activity. If the reverse model is faithful, KL-divergence between the true input and the estimated distributions 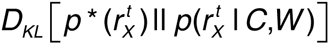 would be minimized (Dayan et al., 1995) (Nessler et al., 2013). Therefore, synaptic weights learning can be performed by 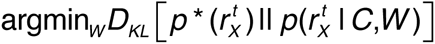. Likelihood 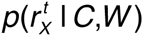 is approximated as

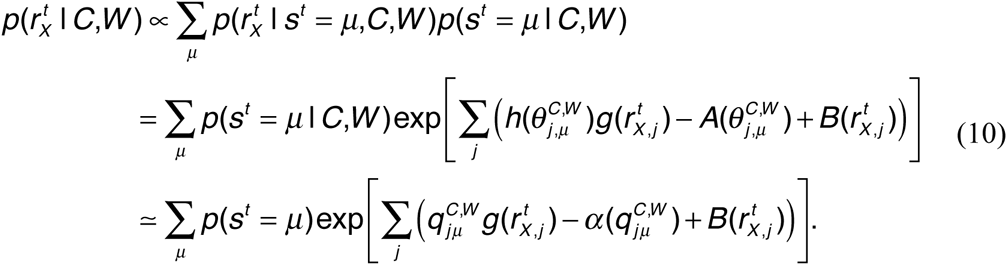

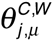 in the second line is the average response estimated from connectivity matrix *C*, and weight matrix *W.* In the last equation, 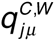 is substituted for 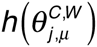. If we approximate the estimated parameter 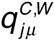 with 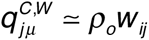 by using the average connectivity *ρ_o_,* a synaptic weight plasticity rule is given by stochastic gradient descending as

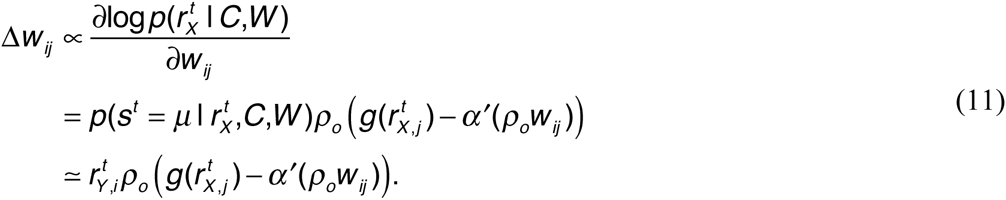

Especially, in a Gaussian model, the synaptic weight converges to the weight coding as 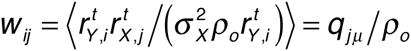, where *μ* is the external state that output neuron *i* learned to represent (i.e. *i* ∈ Ω*_μ_*).

As we were considering population representation, in which the total number of output neuron is larger than the total number of external states (i.e. *p* < *N*), there is a redundancy in representation. Thus, to make use of most of population, homeostatic constraint is necessary. For homeostatic plasticity, we set a constraint on the output firing rate. By combining two terms, synaptic weight plasticity rule is given as

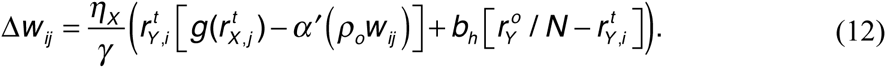

By changing the strength of homeostatic plasticity *b_h_*, the network changes its behavior. The learning rate is divided by *γ*, because the mean of *w* is proportional to 1/*γ*. Although, this learning rule is unsupervised, each output neuron naturally selects an external state in self-organisation manner.

#### 1.5 Synaptic connection learning

Wiring plasticity of synaptic connection can be given in a similar manner. As shown in **Figure 3**, if the synaptic connection structure of network is correlated with the external model, the learning performance typically gets better. Therefore, by considering 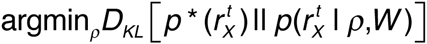, the update rule of connection probability is given as

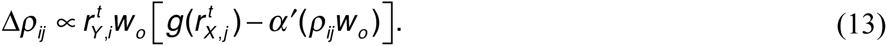

Here, we approximated *w_ij_* with its average value *w_o_*. In this implementation, if synaptic weight is also plastic, convergence of *D_KL_* is no longer guaranteed, yet as shown in **Figure 3**, redundant representation robustly provides a good heuristic solution.

Let us next consider the implementation of the rewiring process with local spine elimination and creation based on the connection probability *ρ_ij_*. To keep the detailed balance of connection probability, creation probability *c_p_* (*ρ*) and elimination probability *e_p_* (*ρ*) need to satisfy

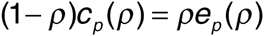

The simplest functions that satisfy above equation is *c_p_* (*ρ*) ≡ *ρ*/*τ_c_*, *e_p_* (*ρ*) ≡ (1–*ρ*) *ρ*/*τ_c_*. In the simulation, we implemented this rule by changing *c_ij_* from 1 to 0 with probability (1–*ρ*)/*τ_c_* for every connection with *c_ij_*=1, and shift *c_ij_* from 0 to 1 with probability *ρ*/*τ_c_* for non-existing connection (*c_ij_*=0) at every time step.

#### 1.6 Dual Hebbian rule and estimated transfer entropy

The results in the main texts suggest that non-random synaptic connection structure can be beneficial either when that increases estimated transfer entropy or is correlated with the structure of the external model. To derive dual Hebbian rule, we used the latter property, yet in the simulation, estimated transfer entropy also increased by the dual Hebbian rule. Here, we consider relationship of two objective functions. Estimation of the external state from the sampled inputs is approximated as

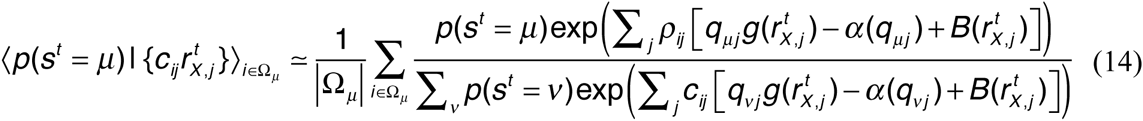

Therefore, by considering stochastic gradient descending, an update rule of *ρ_ij_* is given as

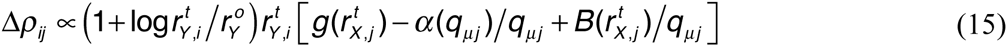

If we compare this equation with the equation for dual Hebbian rule (equation (13)), both of them are monotonically increasing function of 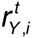 and have the same dependence on 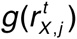 although normalization terms are different. Thus, under an adequate normalization, the inner product of change direction is on average positive. Therefore, although dual Hebbian learning rule does not maximize the estimated maximum transfer entropy, the rule rarely diminishes it.

#### 1.7 Gaussian model

We constructed mean response probabilities 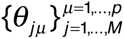 by following 2 steps. First, non-normalized response probabilities 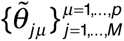 were chosen from a truncated normal distribution *N*(*μ_M_, σ_M_*) defined on [0,∞). Second, we defined 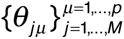 by 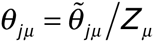, where 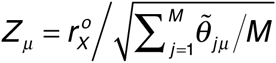. When the noise follows a Gaussian distribution, the response functions in equation (5) are given as

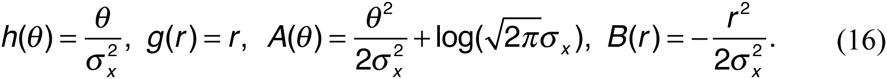

Because 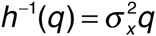, 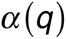 is given as 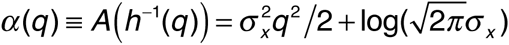. By substituting above values into the original equations, the neural dynamics is given as

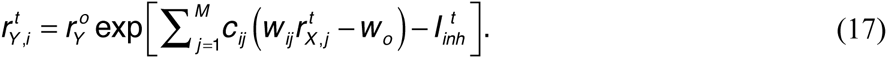

Similarly, dual Hebbian rule becomes

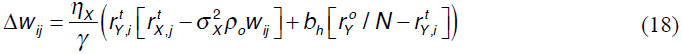

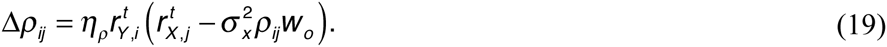

#### 1.8 Poisson model

For Poisson model, we defined mean response probabilities 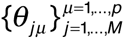 from a log-normal distribution instead of a normal distribution. Non-normalized values were sampled from a truncated log-normal distribution 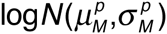 defined on 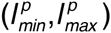. Normalization was performed as 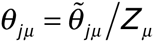 for 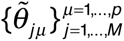, where 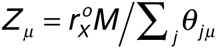. Because the noise follows a Poisson distribution *p*(*r* | *θ*) = exp[−*q + r* log*q* − log*r*!], the response functions are given as

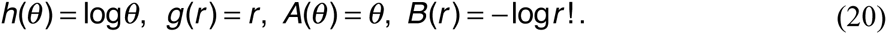

As a result, *α*(*q*) is defined as 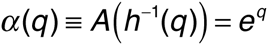. By substituting them to the original equations, the neural dynamics also follows equation (17). If connection is all-to-all, by setting *w_ij_* = log*θ_jμ_*/*θ_o_* for *i* ∈ Ω*_μ_*, optimal inference is achievable. Here, we normalized *θ_j_* by *θ_o_,* which is defined as 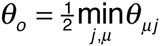, in order to keep synaptic weights in non-negative values.

Learning rules for synaptic weight and connection are given as

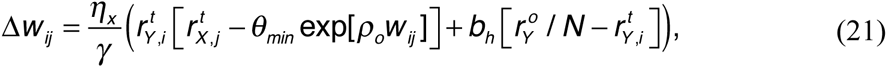

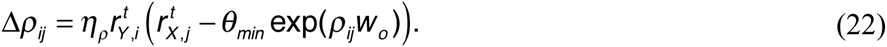

Note that the first term of the synaptic weight learning rule coincides with a previously proposed optimal learning rule for spiking neurons (Nessler et al., 2013)(Habenschuss et al., 2013). In calculation of model error, error was calculated as 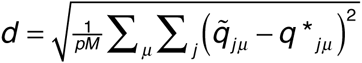, where estimated parameter 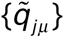 was given by 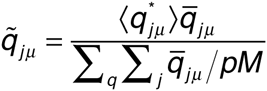. Here, 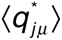 represents the mean of true {*q_jμ_*}, and non-normalized estimator 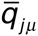 was calculated as 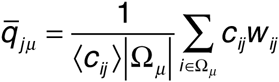. In **Figure S1D**, estimation from connectivity was calculated from 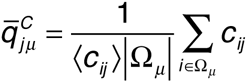, and similarly, estimation from weights was calculated by 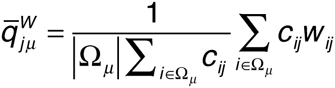. For parameters, we used 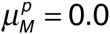, 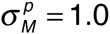, 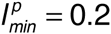, 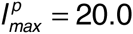, 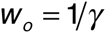, 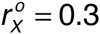, and for other parameters, we used same values with the Gaussian model.

### 2 Analytical evaluations

#### 2.1 Evaluation of performances in weight coding and connectivity coding

In Gaussian model, we can analytically evaluate the performance in two coding schemes. As the dynamics of output neurons follows 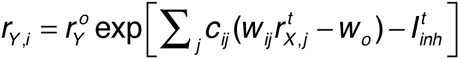, membrane potential variable *u_i_*, which is defined as

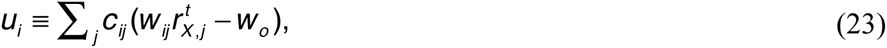

determines firing rates of each neuron. Because {*è_jì_*} is normalized with 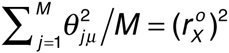, mean and variance of {*è_jì_*} are given as

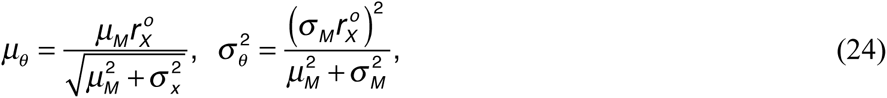

where *μ_M_* and *σ_M_* are the mean and variance of the original non-normalized truncated Gaussian distribution 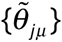. Because both *r_X,j_* and {*è_jì_*} approximately follow Gaussian distribution, *u_i_* is expected to follow Gaussian. Therefore, by evaluating its mean and variance, we can characterize the distribution of *u_i_* for a given external state (Babadi and Sompolinsky, 2014).

Let us first consider the distribution of *u_i_* in the weight coding. In weight coding scheme, *w_jj_* and *c_ij_* are defined as

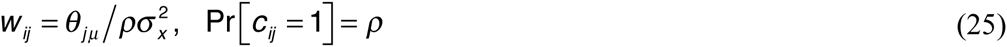

where 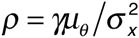. By setting 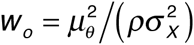, the mean membrane potential of output neuron *i* selective for given signal (i.e. *i* ∈ Ω*_μ_* for *s^t^ = μ*) is calculated as,

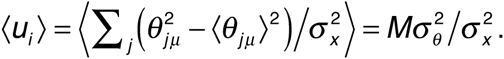

Similarly, the variance of *u_i_* is given as

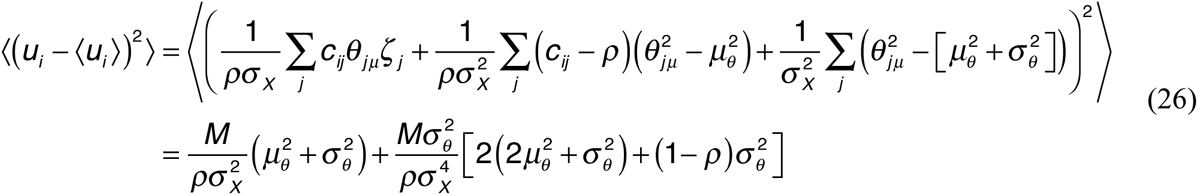

where *ζ_j_* is a Gaussian random variable. On the other hand, if output neuron *i* is not selective for the presented stimuli (if *S^t^* ≠ *μ* and *i* ∈ Ω*_μ_*), *w_ij_* and *r_X,j_* are independent. Thus, the mean and the variance of *u_i_* are given as,

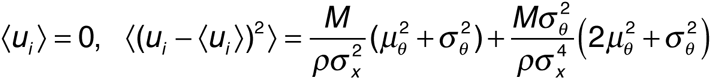

In addition to that, due to feedforward connection, output neurons show noise correlation. For two output neurons *i* and *l* selective for different states (i.e. *i* ∈ Ω*_μ_* and *l* ∉ Ω*_μ_*), the covariance between *u_i_* and *u_l_* satisfies

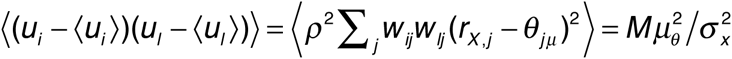

Therefore, approximately (*u_i_*, *u_l_*) follows a multivariable Gaussian distributions

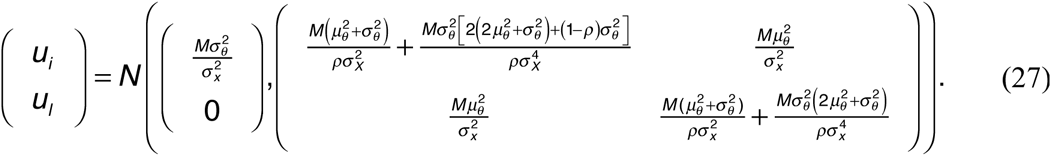

In maximum likelihood estimation, the estimation fails if a non-selective output neuron shows higher firing rate than the selective neuron. When there are two output neurons, probability for such an event is calculated as

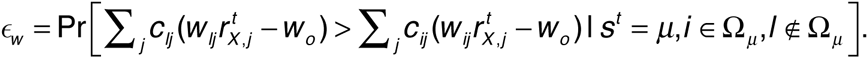

In the simulation, there are *p-1* distractors per one selective output neuron. Thus, approximately, accuracy of estimation was evaluated by (1–*ε_w_*)*^p^*^-1^. In **Figure 2B**, we numerically calculated this value for the analytical estimation.

Similarly, in connectivity coding, *w_ij_* and *c_ij_* are given as

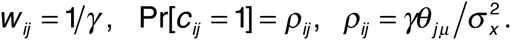

By setting *w_o_ = μ_θ_/γ*, from a similar calculation done above, the mean and the variance of (*u_i_, u_l_*) are derived as

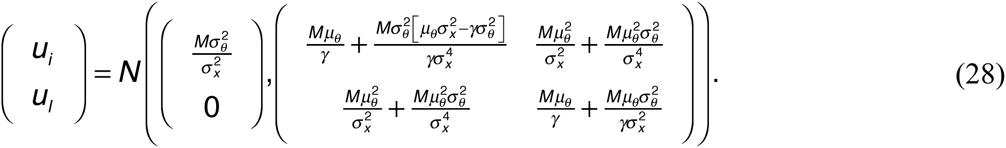

If we compare the two coding schemes, means are the same for two coding schemes, and as *y* satisfies 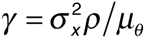, variance of non-selective output neuron are similar. The main difference is the second term of signal variance. In the weight coding, signal variance is proportional to *1/γ,* on the other hands, in the connectivity coding, the second term of signal variance is negative, and does not depend on the connectivity. As a result, in the adequately sparse regime, firing rate variability of selective output neuron becomes smaller in connectivity coding, and the estimation accuracy is better. In the sparse limit, the first term of variance becomes dominant and both schemes do not work well, consequently, the advantage for connectivity coding disappears. Coefficient of variation calculated for signal terms is indeed smaller in connectivity coding scheme (blue and red lines in **Fig 2C**), and the same tendency is observed in simulation (cyan and orange lines in **Fig 2C**).

#### 2.2 Optimality of connectivity

To evaluate optimality of a given connection matrix *C*, we calculated the posterior probability of the external states estimated from *C* and *r_X_*, and compared then to that from the fully connected network *C_all_*. Below, we denote the mean KL-divergence 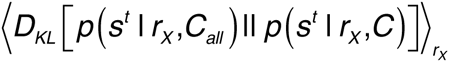 as *I(C_all_, C)* for readability. When the true external state is *s^t^*=*v*, firing rates of input neurons are given by 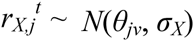, hence this *I(C_all_, C)* is approximately evaluated as

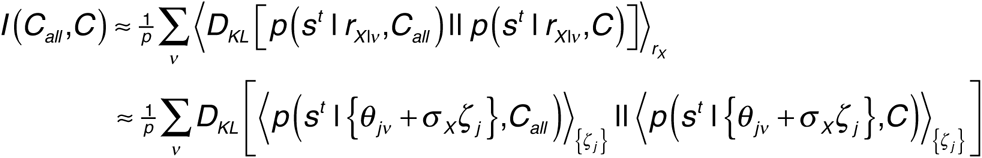

where {*ζ_j_*} are Gaussian random variables, and *C_all_* represents the all-to-all connection matrix. By taking integral over Gaussian variables, the posterior probability is evaluated as

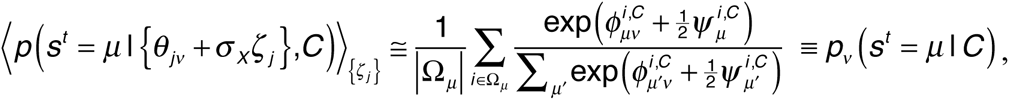

where

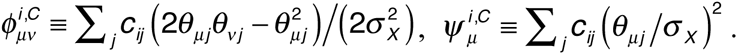

Thus, the KL-divergence between estimations by two connection structures *C_all_* and *C* is approximated as:

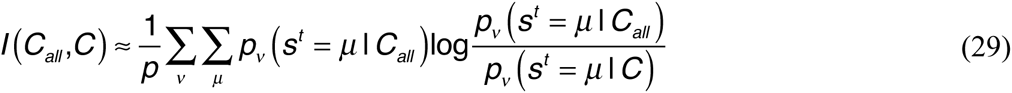

In the black lines in **Figures 3C-E**, we maximized the approximated KL-divergence *I(C_all_,C)* with a hill-climbing method from various initial conditions, thus the lines may not be the exact optimal, but rather lower bounds of the optimal performance. Information gain by a connection *c_ij_* was evaluated by

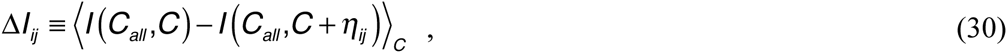

where *η_ij_* is a *N*×*M* matrix in which only (*i*,*j*) element takes 1, and all other elements are 0. In **Figure 3B**, we took average over 1000 random connection structures with connection probability *ρ*=0.1.

### 3 Model settings

#### 3.1 Details of simulation

In the simulation, the external variable *s^t^* was chosen from 10 discrete variables (*p* = 10) with equal probability (Pr[*s^t^* = *q*] = 1/*p*, for all *q*). The mean response probability *θ_jμ_* was given first by randomly chosen parameters 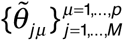 from the truncated normal distribution *N*(*μ_M_,σ_M_*) in [0,∞), and then normalized using 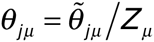, where 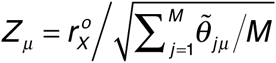. Mean weight *w_o_* was defined as 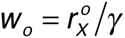. The normalization factor *h_w_* was defined as 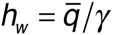 in **Figures 1–2** and **4–5**, where 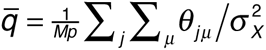, and as 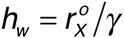 in **Figures 6–7**, as the mean of *θ* depends on *κ_m_.* In **Figure 3**, we used 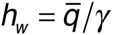 for the dual coding, and 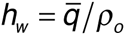 for the rest. Average connectivity 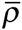 was calculated from the initial connection matrix of each simulation. In the calculation of the dynamics, for the membrane parameter 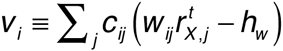, a boundary condition 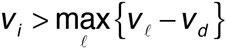 was introduced for numerical convenience, where *v_d_* = -60. In addition, synaptic weight *w_ij_* was bounded to a non-negative value (*w_ij_* > 0), and the connection probability was defined as *ρ* ∈ [0,1]. For simulations with synaptic weight learning, initial weights were defined as 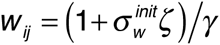, where 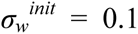, and *ζ* is a Gaussian random variable. Similarly, in the simulation with structural plasticity, the initial condition for the synaptic connection matrix was defined as 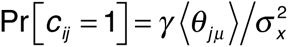. In both the dual Hebbian rule and the approximated dual Hebbian rule, the synaptic weight of a newly created spine was given as 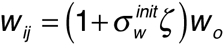, for a random Gaussian variable *ζ* ← *N*(0,1). In **Figure 8**, simulations were initiated at -20 days (i.e., 2 × 10^6^ steps before stimulus onset) to ensure convergence for the control condition. For model parameters, *μ_M_* = 1.0, *σ_M_* = 1.0, *σ_X_* = 1.0, *M* = 200, *N* = 100 *r_X_^o^* = 1.0, and *r_Y_^o^* = 1.0 were used, and for learning-related parameters, *η_X_* = 0.01, *b_h_* = 0.1, *η_ρ_* = 0.001, *τ_c_* = 10^6^, *T_2_* = 10^5^, and *κ_m_* = 0.5 were used. In **Figures 7 and 8**, *η_ρ_* = 0.0001, *τ_c_* = 3 × 10^5^, and *γ* = 0.6 were used, unless otherwise stated.

#### 3.2 Accuracy of estimation

The accuracy was measured with the bootstrap method. By using data from *t-T_o_* <= *t’* < *t*, the selectivity of output neurons was first decided. *Ω_μ_* was defined as a set of output neurons that represents external state *μ.* Neuron *i* belongs to set *Ω_μ_* if *i* satisfies

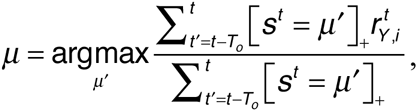

where operator [X]_+_ returns 1 if X is true; otherwise, it returns 0. By using this selectivity, based on data from *t*<= *t’* < *t+T_o_*, the accuracy was estimated as

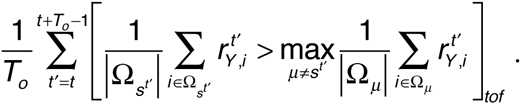

In the simulation, *T_o_* = 10^3^ was used because this value is sufficiently slow compared with weight change but sufficiently long to suppress variability.

#### 3.3. Model error

Using the same procedure, model error was estimated as

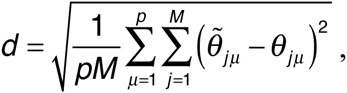

where 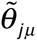 represents the estimated parameter. 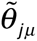 was estimated by

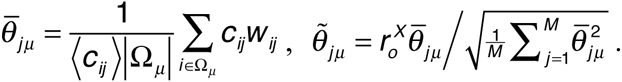

In **Figure 6E**, the estimation of the internal model from connectivity was calculated by

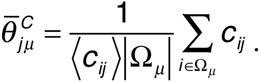

Similarly, the estimation from the synaptic weight in **Figure 6F** was performed with

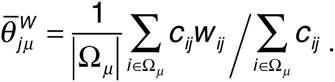

#### 3.4 Transfer entropy

Entropy reduction caused by partial information on input firing rates was evaluated by transfer entropy:

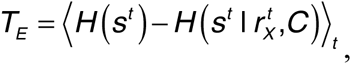

where

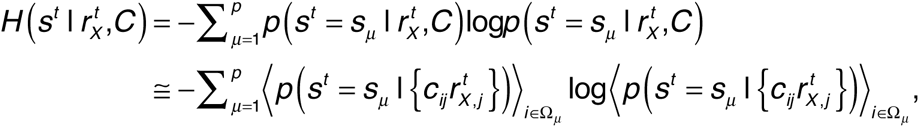

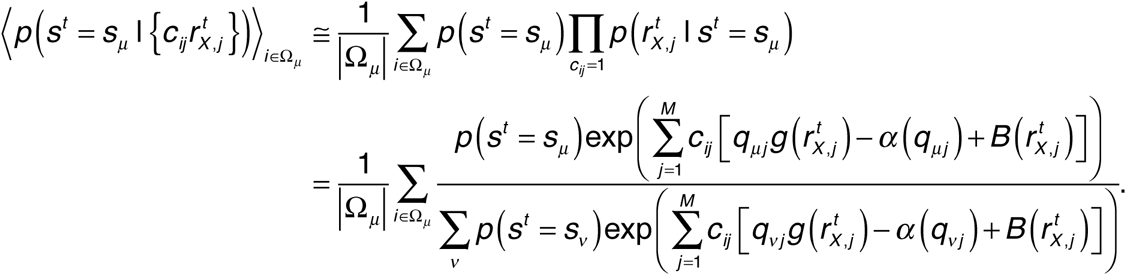

Output group Ω_μ_ was determined as described above. Here, the true model was used instead of the estimated model to evaluate the maximum transfer entropy achieved by the network.

##### Code availability

C++ codes of the simulation program will be available at ModelDB.

## Acknowledgment

The authors thank Drs. Haruo Kasai and Taro Toyoizumi for their comments on the early version of the manuscript.

---

**Supplementary Figure 1:**
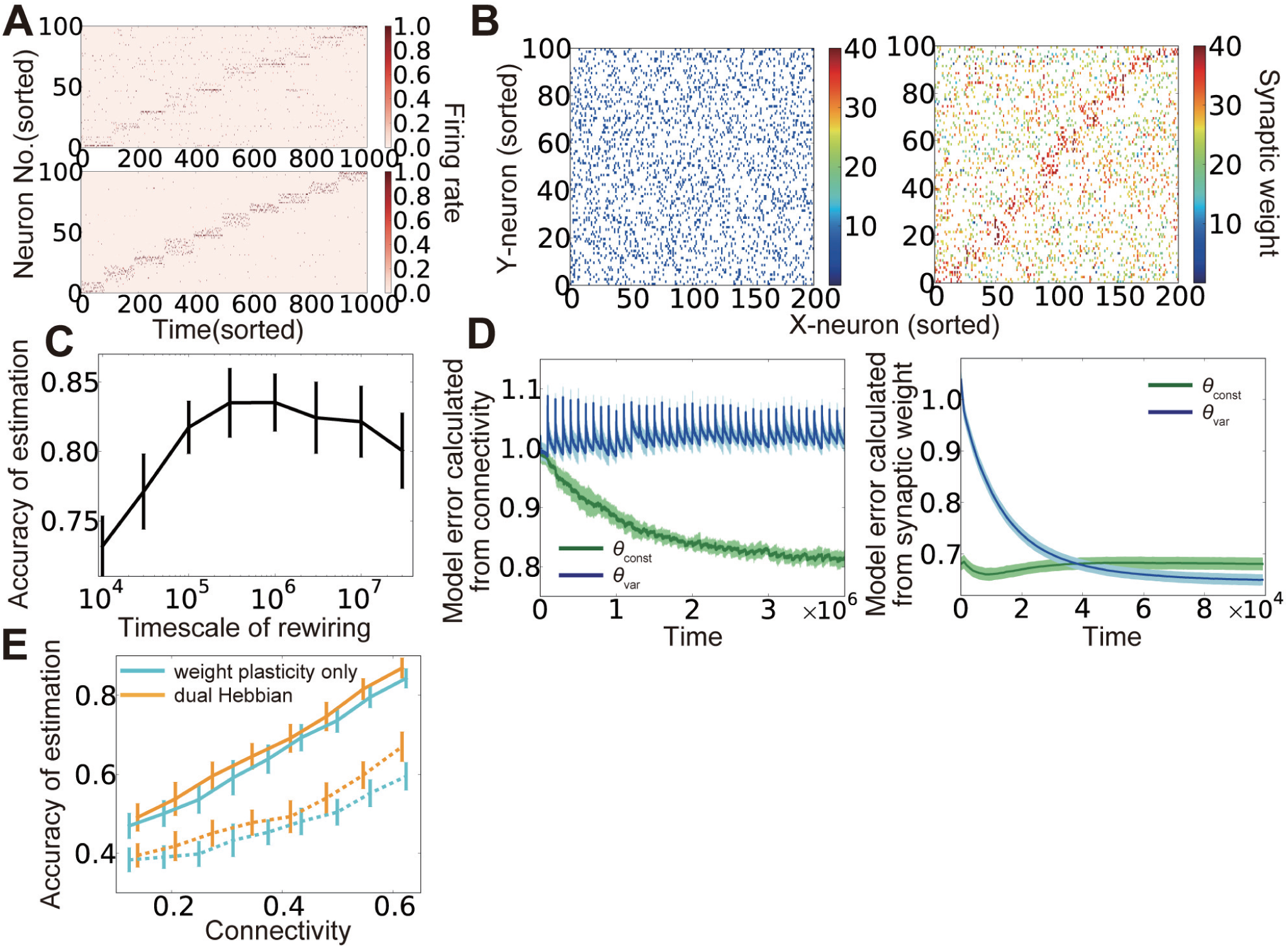
Results in Poisson model. (**A**) An example of output neuron activity before (top) and after (bottom) synaptic weight learning at connectivity *ρ* = 0.25. (**B**) Synaptic weight matrices before (left) and after (right) learning. Both X-neurons and Y-neurons were sorted based on their preferred external states. (**C**) Accuracy of estimation at various timescale of rewiring *τ_c_*. (**D**) Model error calculated from connectivity (left) and synaptic weights (right). (**E**) Comparison of performance among the model without wiring plasticity (cyan), and dual Hebbian model(orange). Corresponding results in the Gaussian model are described in **Fig. 4A**, **Fig. 5F**, **Fig. 5G**, **Fig. 6EF**, **Fig. 6I** respectively.

